# Fitness-driven scaling laws between mRNA and protein levels

**DOI:** 10.1101/2025.03.13.642951

**Authors:** Yichen Yan, Jie Lin

**Affiliations:** Peking-Tsinghua Center for Life Sciences, Peking University, Beijing, China; Center for Quantitative Biology, Peking University, Beijing, China; School of Physics, Peking University, Beijing, China

## Abstract

The connection between protein and mRNA abundance represents a central question in gene expression. While high correlations between mRNA and protein levels are widely observed across different organisms, the evolutionary forces driving these correlations remain elusive. Here, by balancing the cost and benefit of gene expression, we identify the optimal scalings between mRNA and protein levels maximizing cell fitness, where the scaling exponent is positively related to the toxicity effects of protein overexpression. Notably, we predict a lower bound for the exponent, which is 0.5. For the model organism *Saccharomyces cerevisiae*, our model predictions, which incorporate biologically relevant parameters, align well with experimental data. By analyzing genome-wide data across organisms, we find that all organisms exhibit a scaling exponent above the predicted lower bound. These universal phenomena, together with our cost-benefit trade-off model, rationalize the dominance of transcriptional regulation. Our work demonstrates how evolutionary driving forces govern the principles of gene expression regulation.

## Introduction

A cell’s capacity to thrive in a given environment hinges upon its ability to synthesize mRNAs and proteins at the appropriate levels, neither lacking essential components nor wasting resources on superfluous products [1–5]. This intricate balancing necessitates a regulatory system, the principles governing which remain incompletely understood. In this work, we seek to answer a fundamental question: what are the optimal expression levels of mRNAs and proteins that maximize cell fitness? On the one hand, the protein levels should be above some minimum values to perform their functions appropriately [6], and the mRNA levels also cannot be too low; otherwise, the resulting drastic protein noise may compromise the accuracy of gene expression [7–11]. On the other hand, gene expression also imposes a cost on cell fitness, as excessive expression wastes limited cellular resources and disrupts cellular physiological functions [2, 3, 12–18].

To answer the question, we examine the trade-off between the cost and benefit of mRNA and protein production. We present a mathematical model that incorporates the benefit of protein synthesis, the cost associated with transcription and translation, and the toxicity effects of protein overexpression [3]. Numerous experiments suggested that gene expression levels are under evolutionary selection [2, 19–28]. Therefore, we seek to find the optimal mRNA and protein copy numbers that maximize the average fitness. Our numerical simulations and analytical calculations reveal a scaling relationship between the optimal mRNA copy number (*m*) and the protein copy number (*p*): *m* ∼*p*^*γ*^. Surprisingly, the *γ* exponent has a lower bound, 0.5, which occurs when the toxicity effects of protein overexpression are mild; a more substantial toxicity effect results in a larger *γ* exponent.

To verify our theories, we simulate an *in silico S. cerevisiae* cell using biologically relevant parameters. Intriguingly, the predicted *γ* exponent is close to 0.5 for the *in silico* cell, and also agrees well with the experimental data. Furthermore, the predicted scaling between *m* and *p* is observed in diverse organisms [9, 29–32], including human, mouse, fission yeast, and *E. coli*, where the *γ* exponent is all above the predicted lower bound. Particularly, *E. coli* exhibits a scaling close to *m* ∼*p*, suggesting a significant toxicity effect of protein overexpression, consistent with previous experiments [2]. Finally, we reveal the implication of the lower bound of the *γ* exponent: a *γ* exponent larger than 0.5 means that the regulation of protein abundance is dominated by transcription, in agreement with various experiments [30, 33–36]. In summary, our work lays an evolutionary foundation for the mRNA-protein correlation across multiple organisms.

### Benefit and cost of gene expression

We consider a gene whose encoded protein benefits cell fitness but also imposes a cost. We introduce the benefit function *B*(*p*) as the relative increase in fitness due to the produced protein. To simplify without losing the essential ingredient, we set

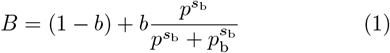

where *b* is a constant. We denote *p*_b_ as the minimal protein number needed, and *s*_b_ is the sensitivity factor quantifying how sensitive cell fitness is to the protein copy number.

We propose that gene expression cost includes two sources. The first source comes from the global competition between genes for finite resources, including ribosome, RNA polymerase [1, 12, 37–44]. Experiments and theories showed that the gene expression load can be decomposed into a transcription cost (*C*_tx_) and a translation cost (*C*_tl_), each proportional to the mRNA copy number and the protein copy number, respectively (Figure 1a) [1, 3, 45–47]. One should note that the competition-generated cost is universal and independent of specific protein functions [47, 48].

**FIG. 1.**
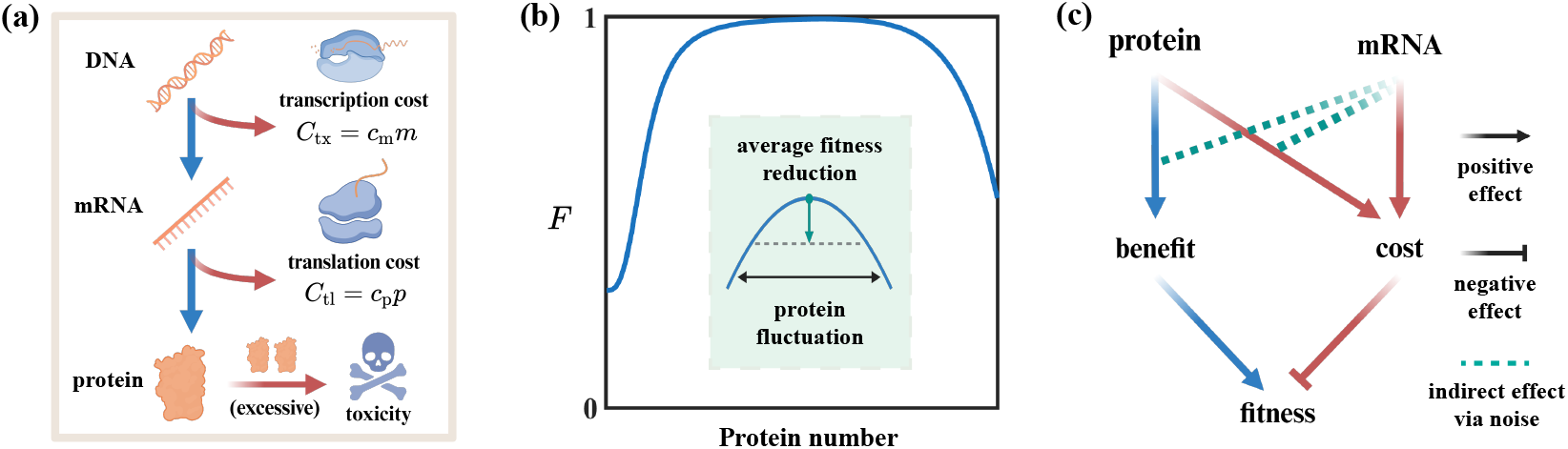
The benefit and cost of gene expression determine the fitness. (a) The cost of gene expression. Gene expression consumes limited cellular resources, imposing a load that reduces cell fitness. The total load lies in both the transcription and the translation processes, each proportional to mRNA and protein copy numbers, respectively. Besides, protein overexpression exacerbates the cost as excessive proteins can be toxic to cells. (b) A schematic of the fitness as a function of protein number. The relative fitness increases from 1 − *b*, approaches 1, and drops with an overexpression of proteins. The inset shows a reduction in the average fitness due to protein noise. (c) The causal maps of how protein and mRNA influence cell fitness. An insufficiency of mRNA gives rise to protein abundance fluctuation, which indirectly affects the benefit and cost of protein expression.

The other source of cost arises from the toxicity of protein overexpression (*C*_d_), which we model by a nonlinear function: 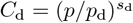. Here, *p*_d_ is a constant which we denote as the toxicity threshold, and *s*_d_ is the sensitivity factor of cell fitness to protein overexpression. To sum up, the total cost function is

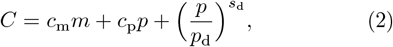

where *c*_m_ is the cost per mRNA copy and *c*_p_ is the cost per protein copy due to resource competition (Figure 1a). Given the cost and benefit, we introduce the total fitness as

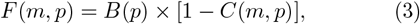

which is a non-monotonic function of protein abundance (Figure 1b), in agreement with previous experiments [19– 21, 23] (Figure S1).

One may expect that the smaller the mRNA copy number, the higher the fitness since it minimizes the transcription cost [Eq. (2)]. However, a small mRNA copy number also leads to a high protein noise, which can reduce the time-averaged or population-averaged fitness (Fig. 1(b)). In this work, we use the following relationship between protein noise and mRNA copy number [49, 50]:

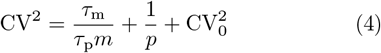

where CV is the coefficient of variation of protein number, *τ*_m_ is the mRNA lifetime, *τ*_p_ is the protein lifetime, and 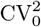 is the noise floor [7, 49, 51–53]. We summarize the cost and benefit of gene expression, where mRNA and protein affect the fitness through coupled pathways (Figure 1c).

### Emerging mRNA-protein scaling

In the following, we find the optimal values of *m* and *p* that maximize the average fitness. For a given gene with its mRNA and protein, we generate 10^5^ random protein numbers following a Gamma distribution with the mean *p* and a variance set by the mRNA copy number *m* according to Eq. (4) [54] and calculate the average fitness ⟨*F* (*m, p*)⟩. We sample a range of mRNA and protein copy numbers to find the *m* and *p* that maximize the average fitness. We change the minimal protein number *p*_b_ to represent proteins with various cellular demands. For simplicity, we keep other parameters the same for all proteins, as the protein copy number needed for physiological functions accounts for the major differences among genes. Intriguingly, a power-law scaling between the optimal mRNA and protein copy numbers emerges (Figure 2a):

**FIG. 2.**
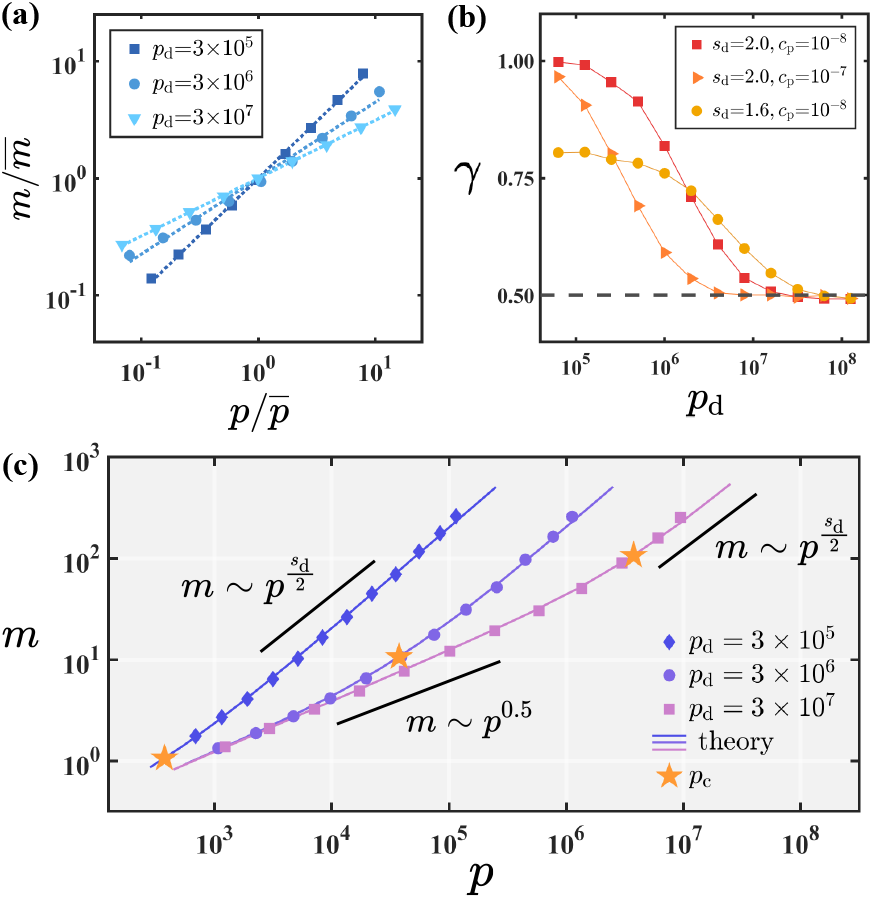
The emerging scaling between mRNA and protein copy numbers at optimal fitness. (a) The mRNA-protein scaling by maximizing the average fitness function considering protein noise, where results with different toxicity thresholds (*p*_d_) are shown. Both mRNA and protein copy numbers are normalized by their mean values. In this figure, we set *τ*_m_*/τ*_p_ = 0.1, CV^2^ = 10^*−*2^, *b* = 0.5, *s*_b_ = 2, *s*_d_ = 2, *c*_m_ = 10^*−*6^, *c*_p_ = 10^*−*8^, and *p*_b_ from 1 to 10^4^ unless otherwise specified. (b) The scaling exponent *γ* decreases with *p*_d_, approaching a lower bound 0.5, regardless of other parameters. (c) The analytical predictions Eq. (7) (solid lines) perfectly align with numerical simulations of the full model. When *p* ≪ *pc, m* ∼ *p*^0.5^, and when *p* ≫ *pc, m* ∼ *p*^*sd/*2^. Here, for each *p*_d_, we scan *p*_b_ from 1 to *p*_d_*/*3.

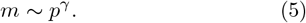

Surprisingly, the *γ* exponent decreases from a value near *s*_*d*_*/*2 and approaches a lower bound 0.5 as the toxicity threshold increases (Figure 2b). In the following, we seek to understand the origin of the emerging scaling theoretically by studying a simplified model, and denote the model mentioned so far as the full model.

### Lower bound of the scaling exponent

We simplify the full model by assuming a small protein noise and approximate the reduction in the average fitness due to protein noise using the local curvature of the fitness function:

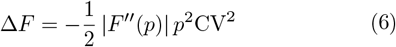

Here, we take *p* at the peak value of the fitness function before considering the noise effects as an approximation; therefore, the first-order derivative does not show up. By further maximizing the fitness with respect to *m* (End Matter), we find the optimal *m* value given the optimal *p* value as

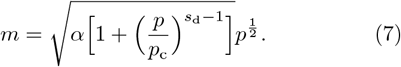

Here, *α* is a constant, and 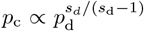 is the critical protein number (see their expressions in End Matter). Intriguingly, Eq. (7) shows that *m* ∼ *p*^0.5^ when the toxicity effect is negligible, i.e., when *p* ≪ *p*_*c*_; it also predicts 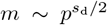 when the toxicity effect is significant, i.e., *p* ≫*p*_*c*_ (Figure 2c).

Intuitively, when toxicity effects are significant, suppressing the protein noise is critical for the cell to survive; therefore, a higher mRNA copy number is necessary to reduce the noise, generating a higher scaling exponent between *m* and *p*. For *p* that is comparable to *p*_c_, the *m* vs. *p* curve exhibits an apparent scaling exponent 0.5 < *γ* < *s*_d_*/*2 (Figure 2c). Indeed, the *m* ∼*p*^*γ*^ scalings from the simulations of the full model (Figure 2a) sit in the different regimes of the analytical prediction of the simplified model (Figure 2c), strongly supporting the validity of the simplified model.

### In silico cells vs. experiments

In the following, we simulate an *in silico* cell using biologically relevant parameters of *S. cerevisiae* (see details in Supplemental Material). In particular, we estimate *s*_b_, *p*_b_, *b, s*_d_, and *p*_d_ by fitting the experimental measured fitness function vs. protein abundance [19]. We also estimate the cost coefficients *c*_*m*_ and *c*_*p*_ based on resource competition [48], supported by experimental measurements [3]. For each gene, we randomly sample parameters for the *in silico* cell from distributions that make good approximations of the experimental data (Table S1 and Figure S2), and compute the optimal mRNA and protein copy numbers, and also the CV^2^ of the protein copy numbers.

We notice that toxicity effects due to protein overex-pression are generally mild in *S. cerevisiae* (Figure S1), suggesting that *p* ≪ *p*_c_ for most genes; therefore, we expect the scaling exponent between *m* and *p* to be close to Indeed, the optimal mRNA and protein copy number exhibits a scaling *m* ∼ *p*^*γ*^ where *γ* ≈ 0.6 (Figure 3a). The CV^2^ of protein number also scales as CV^2^ ∼*p*^*−γ*^ as predicted by Eq. (4), where the upstream noise dominates the protein noise due to a limited mRNA copy number (Figure 3b).

**FIG. 3.**
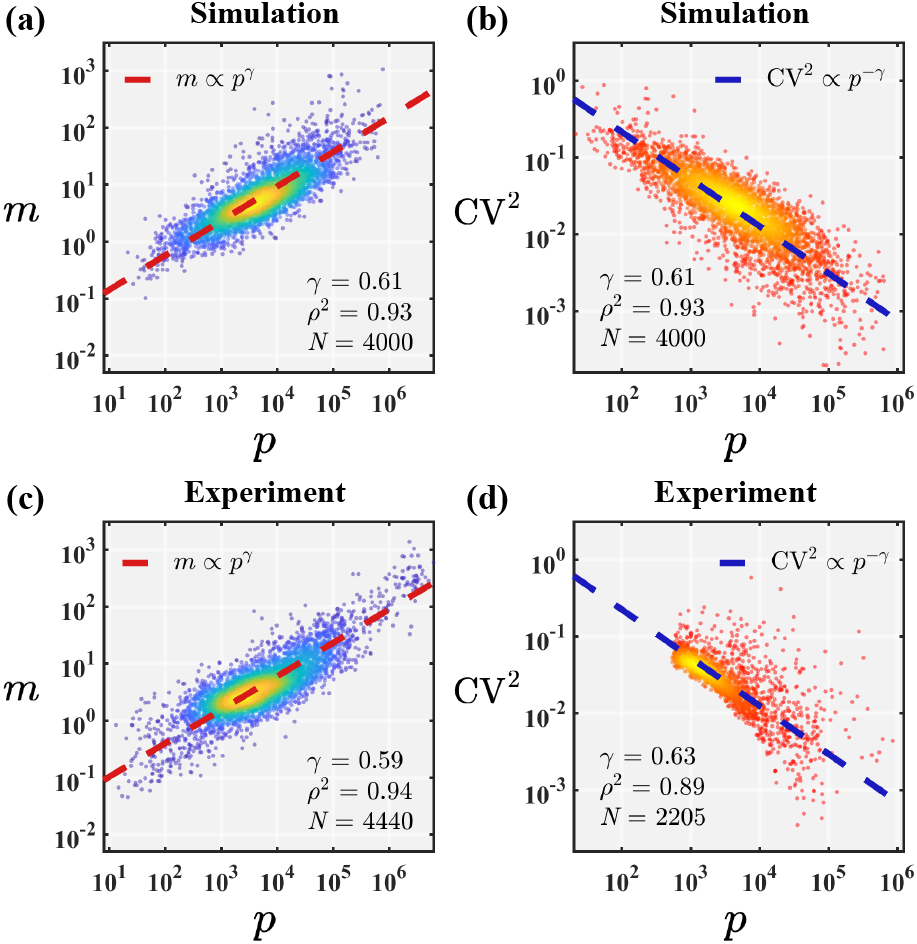
Comparison between the model predictions and experimental data of. *S. cerevisiae*. (a-b) The mRNA copy number and the CV^2^ of protein copy number vs. the protein copy number for the *in silico* cell under the conditions of maximized averaged fitness. The numerical methods are the same as Figure 2. The color of the dots, yellow to blue for (a), red to orange for (b), represents the local density, from high to low. (c-d) The mRNA copy number and the CV^2^ of protein copy number vs. the protein copy number for the experimental data from Ref. [29] and [9], respectively. To clearly show the scaling of CV^2^ vs. *p*, we remove the noise floor 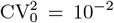 in the plots. The dashed lines (a-d) are power-law fittings. *ρ*^2^ is the fraction of total variance explained by the power-law scaling *m* ∼ *p*^*γ*^, and *N* is the number of genes detected or simulated. The *γ* exponent is calculated by the total-least-squares (TLS) method (Table S2 and Figure S4), which is also used in other relevant figures unless otherwise mentioned.

Surprisingly, the experimental data match our theoretical predictions regarding both the scaling exponent and the absolute values (Figure 3c, d), which strongly supports the validity of our model: balancing the cost and benefit of gene expression is the driving force of gene expression regulation. Meanwhile, the *in silico* simulation demonstrates that our model is robust against genetic heterogeneity since all the genes have different parameters. For *S. cerevisiae*, translational regulation is important because *p/m* ∼ *m*^0.7^: the higher mRNA copy numbers, the more protein copies produced per mRNA copy.

### mRNA-protein scaling across different organisms

We further analyze the genome-wide data of multiple organisms, including *E. coli* [30, 50, 56], *S. cerevisiae* [29, 57], *Schizosaccharomyces pombe* (fission yeast) [31], *Mus musculus* (mouse) [32, 55], and *Homo sapiens* (human) [32, 58]. Intriguingly, all the measured exponents between *m* and *p* are above 0.5 (Figure 4a-d and Figure S3), in perfect agreement with our model predictions. We remark that the lower bound of the *γ* exponent is a consequence of fitness optimization, and no biochemical constraints require that *γ* has to be larger than 0.5.

**FIG. 4.**
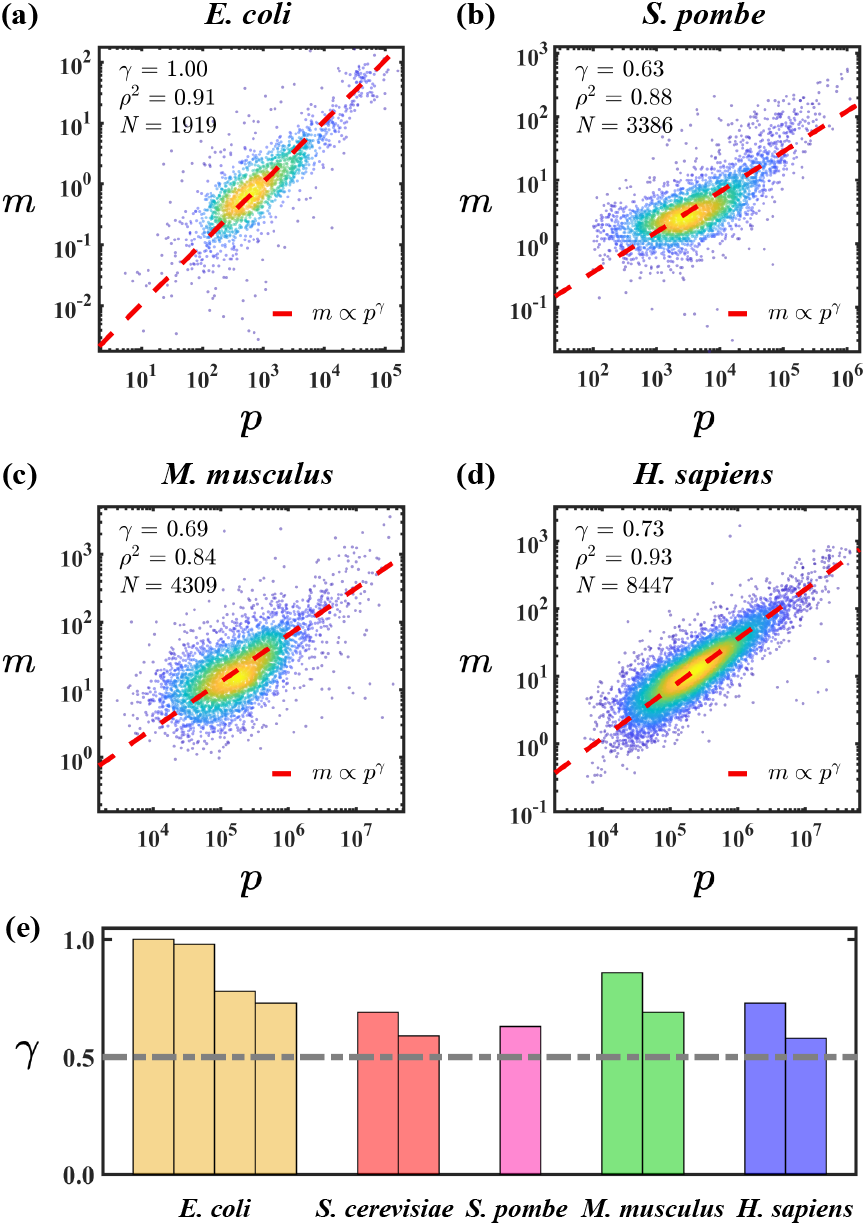
The *m*-*p* scaling across multiple organisms. (a-d) The experimental data of *m* vs. *p* for multiple organisms. The color of the dots (yellow to blue) represents the local density (high to low). Data resources are as follows: *E. coli* [30], *S. pombe* [31], *M. musculus* (NIH 3T3 cell) [55] (the bias in protein measurement corrected by [33]), and *H. sapiens* (HeLa cell) [32]. The color of the dots (yellow to blue) represents the local density (high to low). *ρ*^2^ is the fraction of total variance explained by the power-law scaling *m* ∼ *p*^*γ*^, and *N* is the number of genes detected. (e) The scaling exponents in different species are all higher than 0.5. Each bar represents one dataset, where the original *m*-*p* relations are shown in (a-d) and Figure S3.

What does the lower bound 0.5 mean for the *γ* exponent? Biologically, the expression level of a given protein depends on transcriptional regulation and translational regulation; therefore, one can decompose the protein copy number into two factors *p* = *p*_1_*p*_2_ where *p*_1_ ∼ *m* is the expected protein level if only transcriptional regulation occurs and *p*_2_ represents the extra modulation due to translational regulation. From *m* ∼ *p*^*γ*^, it is straight-forward to find that *p*_2_*/p*_1_ ∼*m*^*δ*^ where 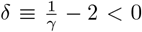. A negative *δ* exponent means that transcriptional regulation dominates over translational regulation, especially for highly expressed genes. Intriguingly, for the model organism *E. coli*, we find a *γ* exponent near 1, suggesting that translational regulation is essentially absent [30].

## Discussion

Optimizing fitness under a cost-benefit trade-off has been applied to determine the expression levels of particular genes both theoretically and experimentally [2, 56, 59, 60]. However, whether a genome-wide scaling between the mRNA and protein expression levels exists due to cost-benefit trade-offs is still elusive. In this work, we unveil a power-law scaling between the mRNA and protein copy numbers that maximizes cell fitness, and predict a lower bound for the *γ* exponent, which is 0.5. We further show that the *γ* exponent depends on the toxicity threshold of protein copy number, and a lower toxicity threshold leads to a higher exponent.

By simulating an *in silico* cell using biologically relevant parameters of *S. cerevisiae*, we demonstrate that our model quantitatively aligns with experiments. In particular, the *γ* exponent for *S. cerevisiae* is near the lower bound 0.5, suggesting a mild toxicity effect of protein overexpression. We further compute the *γ* exponents for multiple organisms and find that in all cases, the exponent is above the predicted lower bound. According to our theory, the different *γ* exponents originate from the distinct toxicity effects of protein overexpression. For *S. cerevisiae*, the expression levels of most proteins are far below the critical protein number (*p* ≪ *p*_c_) such that protein toxicity is neglible; in contrast, for *E. coli*, the toxicity effects are likely to happen near the expression levels of wild-type cells.

We also unveil the implications for the lower bound: protein expression levels are primarily set by mRNA expression levels, in particular for highly expressed genes, in agreement with numerous experiments [30, 33–35, 61– 66]. Our theory does not exclude the possibility of a *γ* exponent larger than 1, and it will be interesting to see whether this situation can happen in some organisms.

## Acknowledgments

The research was funded by the National Key Research and Development Program of China (2024YFA0919600) and supported by Peking-Tsinghua Center for Life Sciences grants.

## End Matter

We first find the optimal *p* that maximizes the fitness before considering the noise effects, where

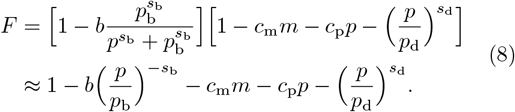

Here, we use the approximation that the reduction in the benefit function relative to its maximum value 1 due to protein deficiency and the cost are both small around the optimal *p*; therefore, higher-order terms are neglected, and the optimal *p* value should be much larger than the minimal protein number *p*_b_ [5]. The agreement between the theoretical predictions and the numerical calculations of the full model supports the validity of these approximations. We compute *∂F* (*m, p*)*/∂p* = 0 and find that

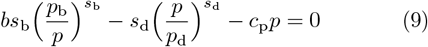

We further calculate the second derivative of the fitness function at the optimal *p* value,

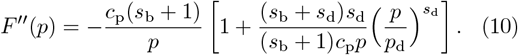

We next find the optimal *m* value by computing *∂*⟨*F* (*m, p*)⟩*/∂m* = 0 where

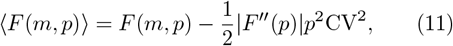

and find that

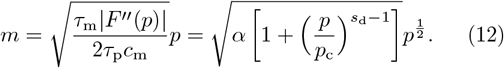

Here,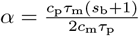, and 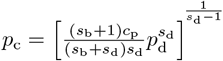.

## Notes

### Competing Interest Statement

The authors have declared no competing interest.

### Summary of Updates

The toxicity effect of protein overexpression is included, which makes the fitness function more biologically relevant.

