## Supplemental Material for "Fitness-driven scaling laws between mRNA and protein levels"

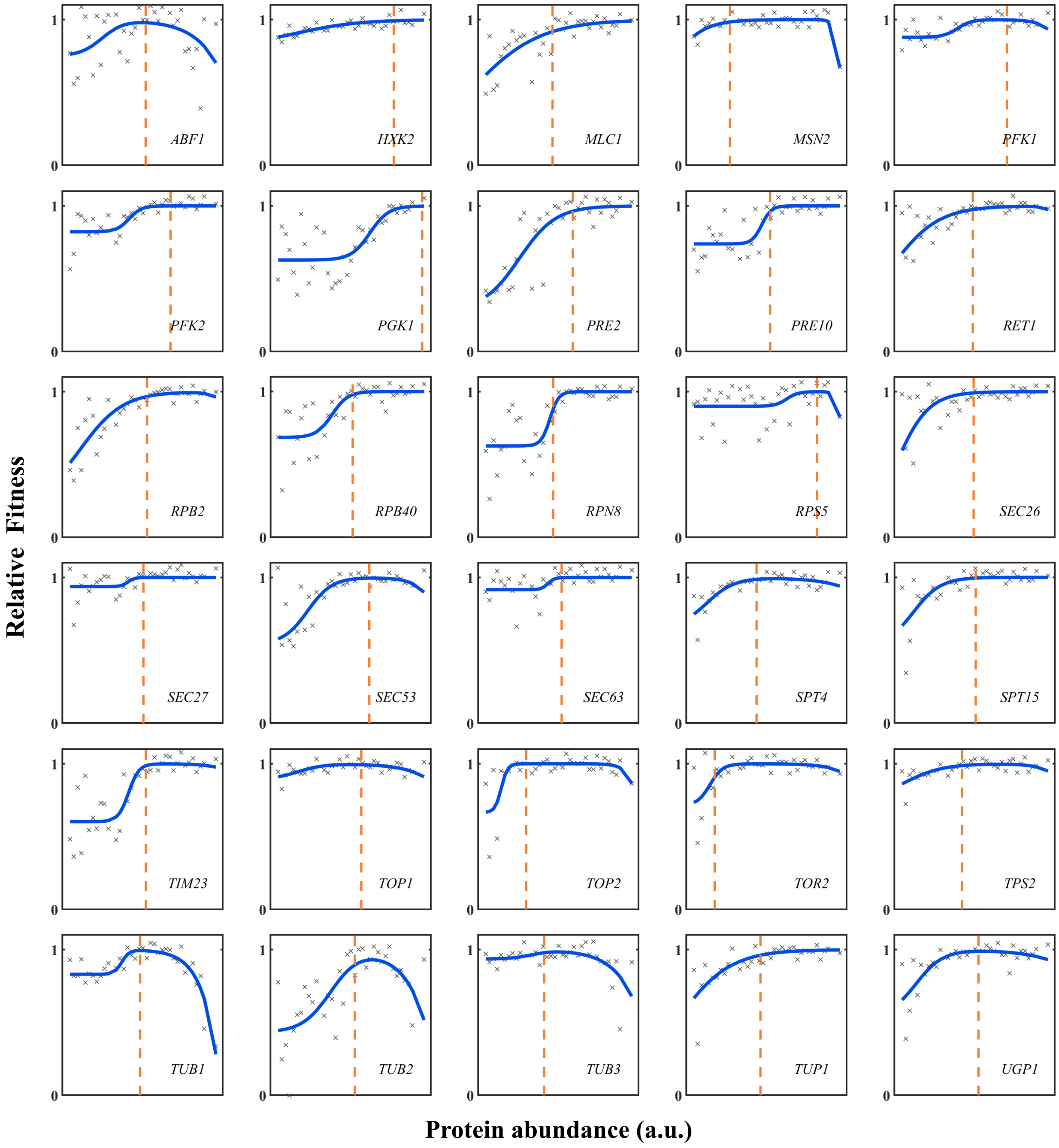


FIG. S1. **The fitness functions in** ***S. cerevisiae*** **for 30 genes.** The crosses are the experimental data from Ref. [1], and the solid line shows the fitting of the fitness function. The orange dashed line marks the wild-type expression level.

### Fitting of the experimental fitness

Keren et al. [1] measured the effects of protein levels on fitness in glucose. We fit the experimental data using the expression of the fitness function, Eq. (3) in the maintext. Here, the transcription cost is neglected since its overall contribution is generally much smaller than the benefit and cost of proteins. $c_{\mathrm{p}}$ is taken from the theoretical calculation in the next section, because it is quite small ($\sim 10^{-8}$), and estimating it from the fittings is presumably inaccurate. We exclude genes that bring no fitness gain (<0.01), leaving 68 genes remaining. We take the conversion factor between the protein number and fluorescence level as 100, which is calibrated using the wild-type expression levels [1]. Figure S1 presents examples of the fitting curves. Figure (S2) shows the corresponding parameter distributions.

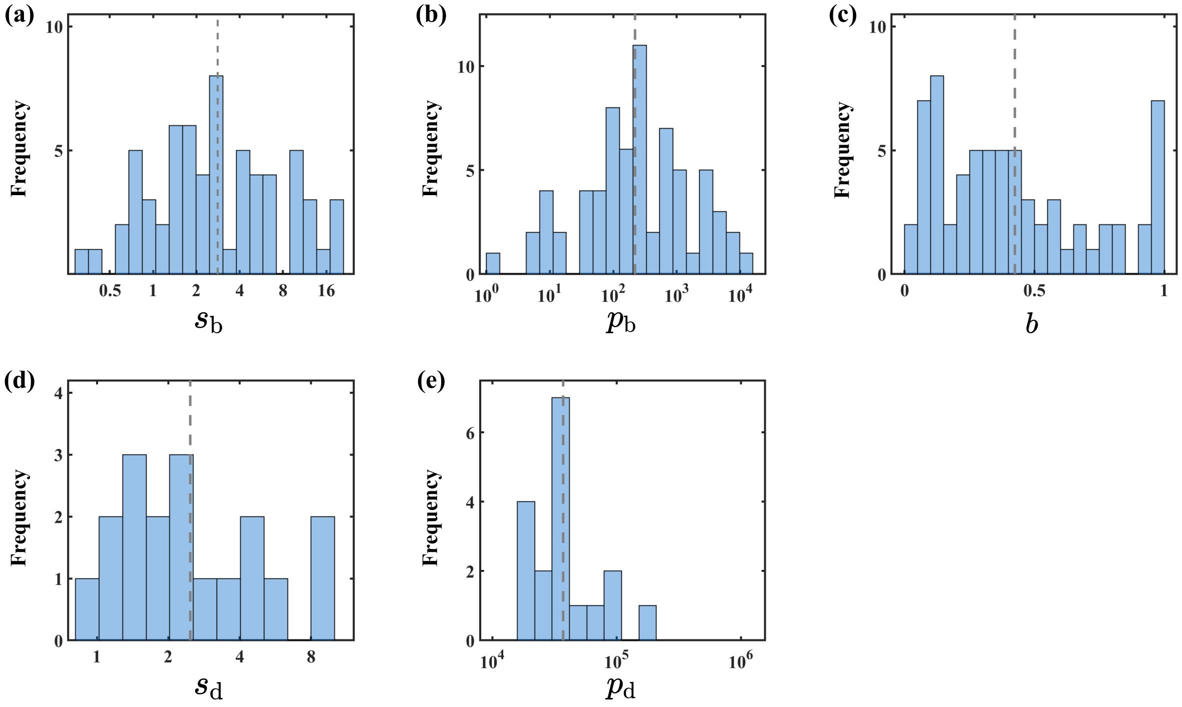


FIG. S2. **The frequency distributions of parameters in the fitness function.** Parameters are extracted from experiments measuring the effects of protein levels on fitness [1]. The dashed line marks the mean value. Within the range of experimental measurements, only 18 genes are detected to be toxic, with $p_d \sim 10^4 - 10^5$.

### Parameters of the *in silico* simulation

We first determine the mean parameter values. We set the average mRNA lifetime as 14 min for *S. cerevisiae*, which agrees with several experimental measurements (12 min [2], 14 min [3], 16 min [4]). The protein lifetime is calculated by $\tau_\mathrm{p} = \frac{1}{\ln 2}/(\frac{1}{1.6\mathrm{h}} + \frac{1}{10\mathrm{h}}) = 2$ h where 1.6 h [5] is the cell doubling time (dilution effect) and 10 h is the average protein degradation half-life [6]. We notice that the protein half-life is mainly set by the cell doubling time [6–8]; therefore, a longer protein degradation half-life [9] has a mild effect on $\tau_\mathrm{p}$.

We take $s_\mathrm{b} = 3$, $s_\mathrm{d} = 3$, and $p_\mathrm{b} = 2 \times 10^2$ according to the results in Figure S2. For $p_\mathrm{d}$, we notice that $p_\mathrm{d} = 10^{4.6}$ for about 1/4 of the genes, and the protein toxicity of the other 3/4 is not detected within a limited expression range. For the latter genes, we estimate the average $p_\mathrm{d}$ to be $10^{6.5}$, which is between the detection limit ($\sim 10^{5.3}$) and the total cellular protein copy number ($\sim 10^{7.7}$ [10, 11]). Averaging the above two parts on a logarithmic scale, the overall average $p_\mathrm{d}$ is estimated to be $1 \times 10^6$.

We use $c_\mathrm{p} = \frac{L_\mathrm{p}}{N_\mathrm{r} v_\mathrm{tl} \tau_\mathrm{p}}$ and $c_\mathrm{m} = \frac{L_\mathrm{m} f_\mathrm{r}}{N_\mathrm{n} v_\mathrm{tx} \tau_\mathrm{m}}$ to estimate the translation cost per protein and the transcription cost per mRNA, which are from our model based on the ribosome and RNAP resource competition [12]. Here, $L_\mathrm{p} = 440$ aa is the protein length [13, 14], $L_\mathrm{m} = 1.5 \times 10^3$ nt is the length of a gene transcribed into mRNA [15, 16], $N_\mathrm{r} = 3 \times 10^5$ is the ribosome copy number [17, 18], $f_\mathrm{r} = 0.3$ is the fraction of inactive ribosomes [19], $N_\mathrm{n} = 3 \times 10^4$ is the RNAP II copy number [20], $v_\mathrm{tl} = 10$ aa/s is the translation elongation speed of ribosome [21, 22], and $v_\mathrm{tx} = 20$ nt/s is the transcription elongation speed of RNAP II [23, 24]. According to the above information, $c_\mathrm{p}$ and $c_\mathrm{m}$ are calculated to be $2 \times 10^{-8}$ and $1 \times 10^{-6}$, respectively. Given a cellular mRNA count of $3 \times 10^4$ molecules and a protein count of $5 \times 10^7$ molecules [10, 11, 25, 26], the total translation cost is approximately 30-fold higher than that of transcription. These results are consistent with theoretical estimations based on ATP consumption rates, where the total translation cost is approximately $10^1$ to $10^2$ times greater than the total transcription cost [15, 27]. Kafri et al. measured the fitness cost of mCherry gene expression, from which we estimate the translation cost per mCherry protein to be $6 \times 10^{-9}$ [5, 12], in agreement with the above calculation given that mCherry is about half the size of an average protein.

To account for genetic heterogeneity, we randomly sample genetic parameters from distributions that provide good approximations of the experimental data. To be specific, $b$ is sampled from a uniform distribution between 0 and 1. Other parameters (denoted by $X$) are sampled from the lognormal distribution $\log_{10} \frac{X}{\langle X \rangle} \sim N(0, \sigma^2)$, where $\langle X \rangle$ represents the above-mentioned mean values, and $\sigma$ is the $\log_{10}$-scale standard deviation. We set $\sigma = 0.3$ for $\tau_\mathrm{m}$ so that 90% of them are within one order of magnitude. The standard deviation of $\tau_\mathrm{p}$ should be small, since it is mainly determined by the cell doubling time, and we set $\sigma = 0.1$ such that 87% of them differ by less than two-fold. The standard deviations of $s_\mathrm{b}$, $s_\mathrm{d}$, $c_\mathrm{m}$, and $c_\mathrm{p}$ are assigned to be 0.15, corresponding to a three-fold range covering about 90% of parameters. The heterogeneity of $c_\mathrm{m}$ and $c_\mathrm{p}$ can originate from differences in gene length, elongation speed, amino acid compositions, etc [12, 28]. Considering the protein demand itself is the major difference among genes, we set $\sigma = 1.0$ for $p_\mathrm{b}$, spanning about four orders of magnitude. The distribution of toxicity threshold is presumably narrower, as proteins typically exhibit toxicity only when present in high abundances, yet cellular tolerance has an upper limit. For instance, cells generally cannot survive if a single protein constitutes most of the proteome. Therefore, we assign a smaller standard deviation $\sigma = 0.4$ for $p_\mathrm{d}$. Alternative parameter choices do not affect our main conclusions.

TABLE S1. **Parameters for the *in silico* simulations of *S. cerevisiae* cell.** $\langle X \rangle$ represents the mean of the distribution, which is extracted from related experiments or theories as mentioned above. All parameters in the following table are sampled from the lognormal distribution $\log_{10} \frac{X}{\langle X \rangle} \sim N(0, \sigma^2)$, i.e., $\log_{10} \frac{X}{\langle X \rangle}$ follows a normal distribution with mean zero and standard deviation $\sigma$. $b$ is sampled from a uniform distribution between 0 and 1.

| Parameter | $c_\mathrm{p}$ | $c_\mathrm{m}$ | $p_\mathrm{b}$ | $p_\mathrm{d}$ | $s_\mathrm{b}$ | $s_\mathrm{d}$ | $\tau_\mathrm{m}$ (min) | $\tau_\mathrm{p}$ (min) |
|---|---|---|---|---|---|---|---|---|
| $\langle X \rangle$ | $2 \times 10^{-8}$ | $1 \times 10^{-6}$ | $2 \times 10^2$ | $1 \times 10^6$ | 3 | 3 | 14 | 120 |
| $\sigma$ | 0.15 | 0.15 | 1.0 | 0.4 | 0.15 | 0.15 | 0.3 | 0.1 |

#### Experimental $m$-$p$ relationships across multiple organisms

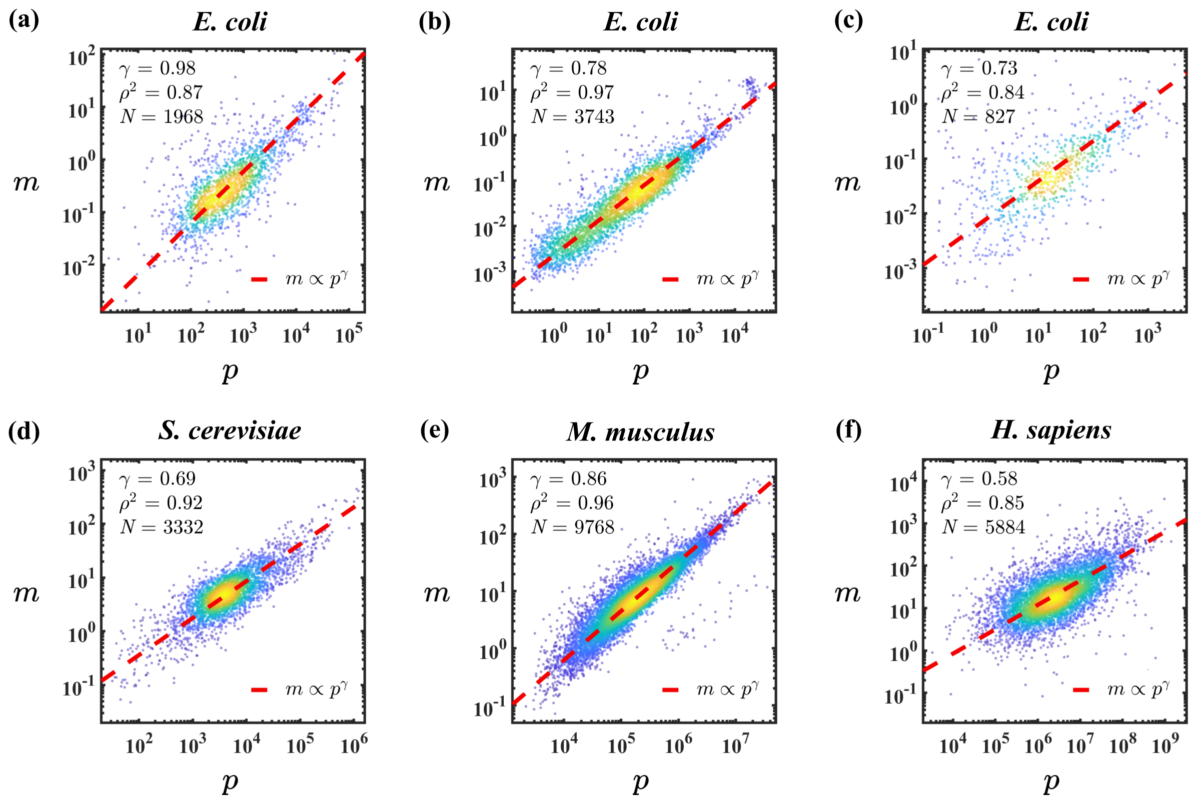


FIG. S3. **The experimental data of $m$ vs. $p$ not included in Figure 4 of the maintext.** The data of (a-f) are from Ref. [29], [30], [31], [32], [33], and [34], respectively. The original results in (d) are presented in number fractions, and here we multiply them by a total mRNA number $3 \times 10^4$ [10, 25] and a total protein number $5 \times 10^7$ [11, 26], respectively, which does not affect the scaling exponent.

### Statistics on the $\gamma$ exponent

The ordinary-least-squares (OLS) linear regression considers only the noise in the vertical-axis-variable and assumes the horizontal-axis-variable is perfectly measured without noise, which is not true in reality [35]. Therefore, the OLS regression fails to capture the $\gamma$ exponent accurately; particularly, the $m$ vs. $p$ slope is not the inverse of the $p$ vs. $m$ slope [36]. In contrast, the total-least-squares (TLS) method accounts for errors or uncertainties in both dimensions and minimizes the sum of the squared perpendicular distances from the data points to the fitted line [37]. Consequently, TLS more faithfully and robustly reflects the relationship between mRNA and protein levels with a stronger explanatory power [38, 39].

Here, we compare the results by different statistical methods (Table S2 and Figure S4). The OLS regression yields a biased lower slope, an effect known as regression dilution bias [40, 41]. Such deviation is especially significant when regressing $p$ on $m$, due to considerable noise in mRNA levels, which may lead to ambiguity regarding the $m$-$p$ correlation. Therefore, all scaling exponents are calculated by the TLS method in this work.

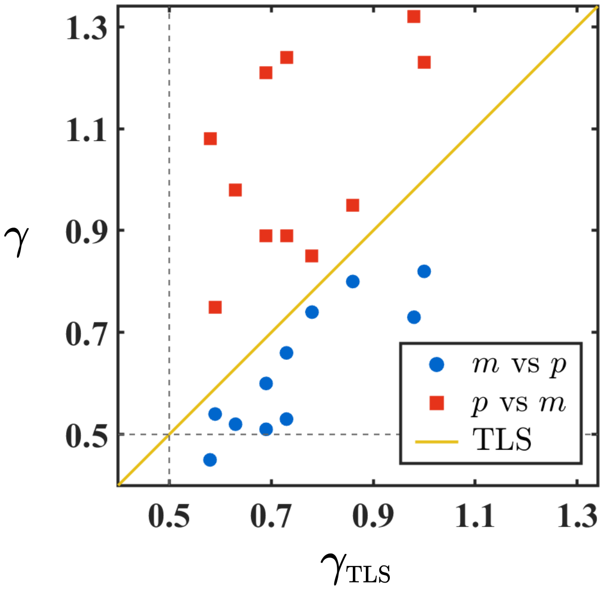


FIG. S4. **The $m$-$p$ scaling exponents $\gamma$ under different statistical methods.** The fitting of $m$ vs. $p$ in the log scale gives smaller scaling exponents than the TLS method, while the $p$ vs. $m$ fitting generates larger $m$-$p$ scaling exponents. The solid line marks $\gamma = \gamma_{\mathrm{TLS}}$ as a reference. The results of $p$ vs. $m$ strongly deviate from the TLS results, indicating considerable noise in mRNA measurements (see also RDR in Table S2). The dashed lines mark the minimum scaling exponent of 0.5, as predicted by our theory, which is not affected by statistical methods. One should notice that in the $p$ vs. $m$ method, $\gamma$ is the inverse of the regression slope. For a more detailed comparison, see Table S2.

TABLE S2. **The statistics on the $m$-$p$ scaling in multiple datasets.** $\gamma$ is the scaling exponent, which is the slope of the regression line; $\rho^2$ is the proportion of variance explained. The TLS method leads to the highest $\rho^2$ in all cases. We introduce the regression dilution ratio (RDR), the ratio between the slope calculated by the standard linear regression divided by the slope of TLS (0<RDR<1). A small RDR indicates large measurement errors of the $x$ variable. We observe that the RDR of the $p$ vs. $m$ method is smaller than that of the $m$ vs. $p$ method, indicating that the measurement errors of mRNA levels are relatively larger than those of protein levels. One should notice that in the $p$ vs. $m$ method, $\gamma$ is the inverse of the regression slope; therefore, the corresponding RDR should be calculated as $\frac{1/\gamma(p \text{ vs. m})}{1/\gamma(\text{TLS})} = \frac{\gamma(\text{TLS})}{\gamma(p \text{ vs. m})}$.

| | Method / Figure | **TLS** | | **$m$ vs. $p$** | | | **$p$ vs. $m$** | | |
|---|---|---|---|---|---|---|---|---|---|
| Species and Data Sources | Fig. | $\gamma$ | $\rho^2$ | $\gamma$ | $\rho^2$ | RDR | $\gamma$ | $\rho^2$ | RDR |
| *E. coli* NCM3722 [29] | 4a | 1.00 | 91% | 0.82 | 67% | 0.82 | 1.23 | 67% | 0.82 |
| *E. coli* NQ390 [29] | S3a | 0.98 | 87% | 0.73 | 55% | 0.75 | 1.32 | 55% | 0.74 |
| *E. coli* MG1655 [30] | S3b | 0.78 | 97% | 0.74 | 86% | 0.95 | 0.85 | 86% | 0.91 |
| *E. coli* DY330 [31] | S3c | 0.73 | 84% | 0.53 | 43% | 0.73 | 1.24 | 43% | 0.59 |
| *S. cerevisiae* BY4741 [33] | 3c | 0.59 | 94% | 0.54 | 72% | 0.91 | 0.75 | 72% | 0.79 |
| *S. cerevisiae* BY4716×BY4700 [32] | S3d | 0.69 | 92% | 0.60 | 68% | 0.88 | 0.89 | 68% | 0.77 |
| *S. pombe* 972 $h^-$ [42] | 4b | 0.63 | 88% | 0.52 | 53% | 0.82 | 0.98 | 53% | 0.64 |
| *M. musculus* NIH 3T3 [43] | S3e | 0.86 | 96% | 0.80 | 84% | 0.93 | 0.95 | 84% | 0.90 |
| *M. musculus* NIH 3T3 [44] | 4c | 0.69 | 84% | 0.51 | 42% | 0.74 | 1.21 | 42% | 0.57 |
| *H. sapiens* HeLa [43] | 4d | 0.73 | 93% | 0.66 | 74% | 0.90 | 0.89 | 74% | 0.82 |
| *H. sapiens* HeLa [34] | S3f | 0.58 | 85% | 0.45 | 41% | 0.78 | 1.08 | 41% | 0.53 |

[8] R. Christiano, H. Arlt, S. Kabatnik, N. Mejhert, Z. W. Lai, R. V. Farese, and T. C. Walther, A systematic protein turnover

map for decoding protein degradation, Cell reports **33** (2020).

[9] J. M. Pratt, J. Petty, I. Riba-Garcia, D. H. Robertson, S. J. Gaskell, S. G. Oliver, and R. J. Beynon, Dynamics of protein turnover, a missing dimension in proteomics, Molecular & Cellular Proteomics **1**, 579 (2002).

[10] F. Miura, N. Kawaguchi, M. Yoshida, C. Uematsu, K. Kito, Y. Sakaki, and T. Ito, Absolute quantification of the budding yeast transcriptome by means of competitive pcr between genomic and complementary dnas, BMC genomics **9**, 574 (2008).

[11] B. Ho, A. Baryshnikova, and G. W. Brown, Unification of protein abundance datasets yields a quantitative saccharomyces cerevisiae proteome, Cell Systems **6**, 192 (2018).

[12] Y. Yan and J. Lin, Disentangling the cost of gene expression, bioRxiv , 2025 (2025).

[13] H. F. Lodish, *Molecular Cell biology* (Macmillan, 2008).

[39] S. Van Huffel and P. Lemmerling, *Total least squares and errors-in-variables modeling: analysis, algorithms and applications* (Springer Science & Business Media, 2013).

[40] N. Draper, *Applied regression analysis* (McGraw-Hill. Inc, 1998).

[41] S. Weisberg, *Applied linear regression*, Vol. 528 (John Wiley & Sons, 2005).

[42] S. Marguerat, A. Schmidt, S. Codlin, W. Chen, R. Aebersold, and J. Bähler, Quantitative analysis of fission yeast transcriptomes and proteomes in proliferating and quiescent cells, Cell **151**, 671 (2012).

[43] S. W. Eichhorn, H. Guo, S. E. McGeary, R. A. Rodriguez-Mias, C. Shin, D. Baek, S.-h. Hsu, K. Ghoshal, J. Villén, and D. P. Bartel, mrna destabilization is the dominant effect of mammalian micrornas by the time substantial repression ensues,

Molecular Cell **56**, 104 (2014).
[44] B. Schwanhäusser, D. Busse, N. Li, G. Dittmar, J. Schuchhardt, J. Wolf, W. Chen, and M. Selbach, Global quantification of mammalian gene expression control, Nature **473**, 337 (2011).
